# Host cell neddylation facilitates alphaherpesvirus entry in a virus-specific and cell-dependent manner

**DOI:** 10.1101/2022.07.20.500907

**Authors:** Becky H. Lee, Giulia Tebaldi, Suzanne M. Pritchard, Anthony V. Nicola

## Abstract

Herpes simplex virus 1 (HSV-1) commandeers the host cell proteasome at several steps of its replication cycle, including entry. Here we demonstrate that HSV-2, pseudorabies virus (PRV), and bovine herpesvirus 1 (BoHV-1) entry are blocked by bortezomib, a proteasome inhibitor that is a FDA-approved cancer drug. Proteasome-dependent entry of HSV-1 is thought to be ubiquitin-independent. To interrogate further the proteasomal mechanism of entry, we determined the involvement of the ubiquitin-like molecule NEDD8 and the neddylation cascade in alphaherpesvirus entry and infection. MLN4924 is a small-molecule inhibitor of neddylation that binds directly to the NEDD8-activating enzyme. Cell treatment with MLN4924 inhibited plaque formation and infectivity by HSV-1, PRV and BoHV-1 at non-cytotoxic concentrations. Thus, the neddylation pathway is broadly important for alphaherpesvirus infection. However, the neddylation inhibitor had little effect on entry of the veterinary viruses but had a significant inhibitory effect on entry of HSV-1 and HSV-2 into seven different cell types. Wash-out experiments indicated that MLN4924’s effect on viral entry was reversible. Time-of-addition assay suggested that the drug was acting on an early step in the entry process. siRNA knockdown of NEDD8 significantly inhibited HSV entry. In probing the neddylation-dependent step in entry, we found that MLN4924 dramatically blocked endocytic uptake of HSV from the plasma membrane by >90%. In contrast, the rate of HSV entry into cells that support direct fusion of HSV with the cell surface was unaffected by MLN4924. Interestingly, proteasome activity was less important for the endocytic internalization of HSV from the cell surface. The results suggest that the NEDD8 cascade is critical for the internalization step of HSV entry.

**IMPORTANCE:** Alphaherpesviruses are ubiquitous pathogens of humans and veterinary species that cause lifelong latent infections and significant morbidity and mortality. Host cell neddylation is important for cell homeostasis and for the infection of many viruses, including HSV-1, HSV-2, PRV, and BoHV-1. Inhibition of neddylation by a pharmacologic inhibitor or siRNA blocked HSV infection at the entry step. Specifically, the NEDD8 pathway was critically important for HSV-1 internalization from the cell surface by an endocytosis mechanism. The results expand our limited understanding of cellular processes that mediate HSV internalization. To our knowledge, this is the first demonstration of a function for the neddylation cascade in virus entry.

## INTRODUCTION

Alphaherpesviruses are ubiquitous pathogens that are host-adapted to many mammalian species. Some members of this herpesvirus subfamily have a propensity to infect mucosal epithelial cells as a portal of entry into the host and establish latency in the peripheral nervous system. Immunocompetent individuals infected by alphaherpesviruses often manifest little to no clinical disease. Neonates or immunocompromised individuals present with debilitating diseases and sometimes fatal infection. Herpes simplex virus 1 (HSV-1), the prototype human alphaherpesvirus, most commonly causes cold sores but can lead to blindness, encephalitis, or fatal neonatal infections (1–3). An effective HSV vaccine that prevents infection remains elusive (4–6). HSV-2 is the most common cause of genital ulcers in humans, and genital herpes infection increases the acquisition and transmission of HIV infection (7, 8). The porcine alphaherpesvirus, pseudorabies virus (PRV) most commonly causes neurologic disorders, reproductive failure, and poor growth rate in production pigs, which leads to significant losses in the swine industry worldwide (9, 10). Bovine herpesvirus 1 (BoHV-1) most commonly causes severe ulcerative rhinotracheitis in cattle and is a significant contributor to the economically impactful polymicrobial infection known as bovine respiratory disease complex (11–14). Critical aspects of the entry of the alphaherpesviruses studied here are conserved, including nectin receptor usage and dependence on host cell proteasome activity (15–20)

Alphaherpesviruses enter host cells in a cell type-dependent manner (18, 19, 21–26). HSV-1 is internalized by endocytosis into epithelial cells and transits a nonconventional endocytic pathway that requires low pH (27–29). HSV-1 enters neurons by non-endocytic, pH-independent penetration at the plasma membrane. Regardless of cell type, entry of HSV is facilitated by host cell proteasome activity, including the targeting of incoming capsids to the nuclear periphery (17, 20). Transport of entering HSV capsids is considered a post-fusion step in the entry process and is facilitated by the viral E3 ubiquitin ligase ICP0 present in the inner tegument layer (30, 31). HSV enters cells in the absence of a functional host ubiquitin-activating enzyme, suggesting that viral entry is ubiquitin-independent (17). The function of ubiquitin-like moieties, such as neural precursor cell expressed developmentally down-regulated protein 8 (NEDD8), in alphaherpesvirus entry and infection are incompletely understood and is a focus of the current report.

The 26S proteasome is a proteolytic machine that maintains cellular homeostasis primarily through the ubiquitin-proteasome system (UPS) (32–34). Many viruses utilize the 26S proteasome for different steps in the entry process (35). Alphaherpesviruses interact with the UPS to promote different aspects of the viral replicative cycle. See references (17-20, 36, 37) as examples. Inhibition of neddylation has broad antiviral activity (38, 39) but the specific involvement of neddylation in viral entry has not been addressed to our knowledge. Neddylation is the reversible conjugation of the 81 amino acid NEDD8 to a substrate (40–42). A well-described function of neddylation is the activation of cullin RING ligases (CRLs), the largest class of ubiquitin E3 ligases (43). Thus, neddylation can be functionally linked to the UPS and assist in protein degradation. The neddylation cascade may function in regulation of protein stability, alter subcellular localization of proteins, and influence protein-protein interactions (44). Additionally, viruses can modulate CRLs (45) and some rely on host cell neddylation for replication (46–51). We report that neddylation, along with proteasomal degradation, is important for infection by the alphaherpesviruses HSV-1, HSV-2, PRV, and BoHV-1. The NEDD8 cascade facilitated HSV entry, particularly at the level of endocytic internalization of viral particles from the plasma membrane. Interestingly, the internalization step in HSV entry was less dependent on proteasome activity.

## RESULTS

### Alphaherpesvirus entry by a conserved proteasome-dependent mechanism

HSV-1 relies on the cellular proteasome for entry, specifically at a post-penetration step (17). Entry of PRV into PK15 cells and of BoHV-1 into MDBK cells is also proteasome-dependent as it is inhibited by the widely used proteasome inhibitor MG132 (18, 19). To further assess alphaherpesvirus dependence on the proteasome for entry, we tested the effect of bortezomib on entry of HSV-2, BoHV-1, and PRV as measured by β-galactosidase reporter assay (Fig. 1 A-C). Bortezomib is a peptide boronate drug that causes potent, selective, and reversible inhibition of the proteasome by binding directly to the beta-5 subunit of the 20S core of the 26S proteasome (52–54). Bortezomib inhibits HSV-1 at early stages of infection (20). Here we show that bortezomib inhibited HSV-2, PRV, and BoHV-1 entry in a concentration-dependent manner. Bortezomib concentrations that resulted in at least 50% inhibition were 0.075 µM for HSV-2 G (Fig. 1A), 1 µM for PRV BeBlue (Fig. 1B), and 0.1 µM for BoHV-1 v4a (Fig. 1C). Therefore, bortezomib impairs alphaherpesvirus entry, and the inhibitory concentrations were not cytotoxic as determined by lactate dehydrogenase (LDH) assay (Fig 1A-C). Thus, the host cell proteasome activity facilitates entry of several members of the alphaherpesvirus subfamily (17–19).

**Figure 1.**
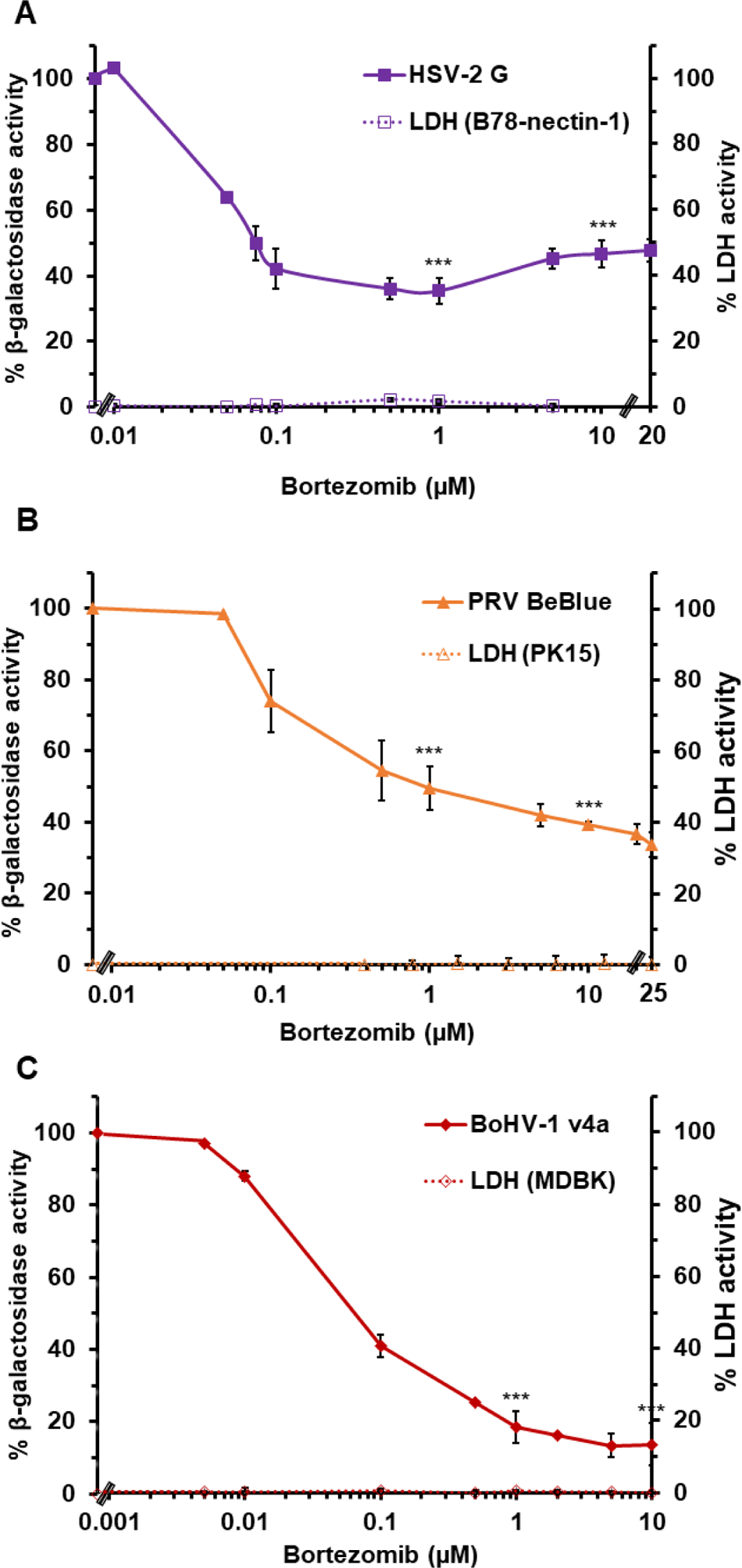
Effect of bortezomib, a proteasome inhibitor, on alphaherpesvirus entry. (A-C) B78-nectin-1, PK15, or MDBK cells were treated with bortezomib for 20 min at 37°C. HSV-2 G (MOI 0.1), PRV BeBlue (MOI 1), or BoHV-1 v4a (MOI 2) was added to the respective cell lines in the continued presence of bortezomib. At 6 h p.i., cell lysates were prepared, and β-galactosidase activity of mock-treated, infected cells was set to 100%. Cytotoxicity is shown as percent LDH activity. Each experiment was performed with quadruplicate (entry) or triplicate (LDH) samples. Values are the means and standard errors of data from three independent experiments. Student’s *t*-test comparing no-drug to either 1 µM or 10 µM bortezomib. (***, P < 0.005)

### Blocking neddylation impairs alphaherpesvirus plaque formation and viral titer

HSV-1 entry does not require an active E1 ubiquitin-activating enzyme; thus, proteasome-mediated entry is thought to be ubiquitin-independent (17). We investigated the role of neddylation, a ubiquitin-like modification, in the proteasome-dependent entry of alphaherpesviruses. MLN4924, also known as pevonedistat, is a small-molecule inhibitor of neddylation (55, 56). It binds to the adenylation site of the NEDD8 activating enzyme (NAE) and interferes with NEDD8 conjugation to substrate proteins (56). MLN4924 has been tested in pre-clinical trials to treat hematologic malignancies (57, 58). MLN4924 has antiviral activity against several viruses including HSV and HCMV (38). We tested the effect of MLN4924 treatment on alphaherpesvirus plaque formation and on the production of infectious progeny virions. Treatment of cells with MLN4924 inhibited plaque formation of HSV-1 on Vero cells (Fig. 2A), PRV on PK15 cells (Fig. 2B), and BoHV-1 on MDBK cells (Fig. 2C) in a concentration-dependent manner. MLN4924 treatment similarly inhibited viral progeny production for HSV-1 (Fig. 2D), PRV (Fig. 2E), and BoHV-1 (Fig. 2F) in a concentration-dependent manner. The concentrations of MLN4924 used in these experiments were not cytotoxic for each of the cell types (Fig. 2G). Up to 40 µM MLN4924 treatment of Vero or PK15 cells had little effect on cell viability as measured by LDH assay (Fig. 2G). Up to 1 µM MLN4924 treatment of MDBK cells yielded little to no effect on viability (Fig. 2G). However, treatment with 2 to 40 µM MLN4924 resulted in MDBK cell cytotoxicity ranging from 24 to 55% (data not shown). A range of lower concentrations of MLN4924 was necessarily used for the BoHV-1 experiments on MDBK cells (Fig. 2C and 2F). All in all, our results suggest that pharmacologic inhibition of host cell neddylation reduces plaque formation and production of viral progeny for the alphaherpesviruses tested.

**Figure 2.**
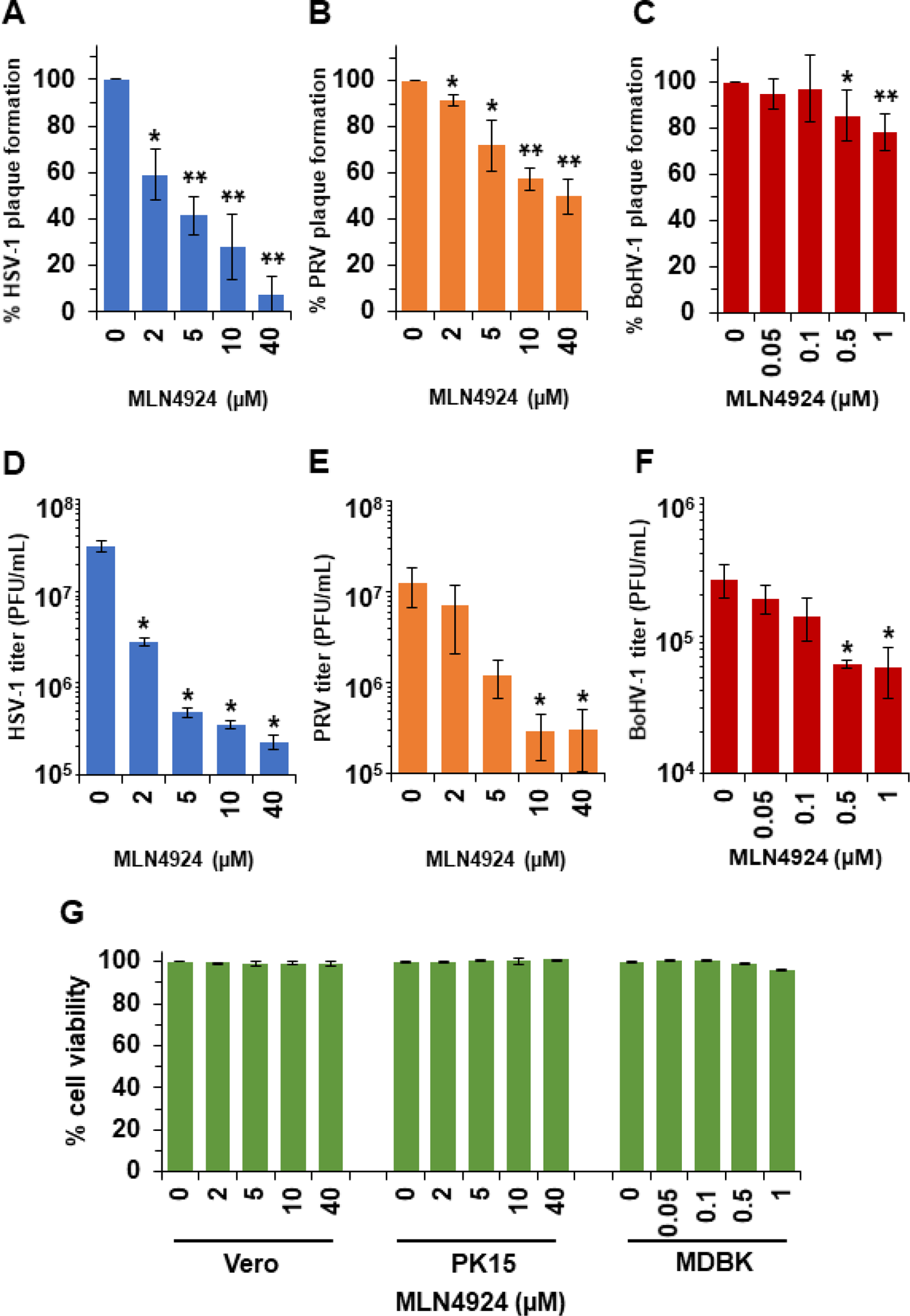
Effect of neddylation inhibitor MLN4924 on alphaherpesvirus plaque formation and infectivity. Vero cells (A, D), PK15 cells (B, E), or MDBK cells (C, F) were treated with MLN4924 for 20 min at 37°C. HSV-1 KOS tk12 (A, D), PRV BeBlue (B, E), or BoHV-1 v4a (C, F) was added in the continued presence of MLN4924. (A-C) Approximately 100 PFU/well was added. At 1-1.5 h p.i., a 1% CMC overlay was added. At 42 to 48 h p.i., cells were fixed, and plaques were enumerated. Plaque number in the no-drug wells was set to 100%. Data are the mean of three independent experiments with standard deviation. (D-F) MOIs were 0.01, 0.0002, or 0.002 for HSV-1 KOS tk12, PRV BeBlue, or BoHV-1 v4a, respectively. At 24 h p.i., cells were scraped from the wells and collected along with the supernatant. This sample, containing cells and supernatant, were subjected to a freeze-thaw cycle and sonication and then titered on the appropriate cell type. Data are the mean of three independent experiments with standard error. (G) MLN4924 exhibits minimal cytotoxicity at drug concentrations used in infectivity experiments. MLN4924 was added to Vero, PK15, or MDBK cells. At 24 h, LDH activity in the supernatant was assayed. Values are percentages of mock-treated (DMSO). Student’s *t*-test (A-C) comparing each indicated MLN4924-treated sample to mock-treated. Wilcoxon Rank Sum Test (Mann Whitney U test) (D-F) comparing MLN4924-treated conditions (2, 5, 10, or 40 µM) to mock-treated (*, P < 0.05; **, P < 0.01).

### Effect of a neddylation inhibitor on alphaherpesvirus entry

We determined whether neddylation functioned during the entry stage of alphaherpesvirus infection. Alphaherpesvirus entry is proteasome-dependent. Thus, we compared the effects of MLN4924 and MG132, a peptide aldehyde drug that inhibits the proteasome, using the β-galactosidase reporter assay for entry (Fig. 3 A-D), which is standard in the field (59–63). The *lacZ^+^* strains of HSV, PRV and BoHV-1 have been employed in many previous studies of viral entry (examples include (15, 18, 19, 64–69). Since reporter gene activity is measured at 6 h p.i., it is important to note that steps up to and including gene expression mediated by the respective promoters might be affected in this assay. MLN4924 inhibited entry of HSV-1 KOS tk12 (Fig. 3A) and HSV-2 G (Fig. 3B) by more than 60%. As expected, MG132 also inhibited HSV-1 and HSV-2 entry in a concentration dependent manner (Fig. 3A, B). MLN4924 was less effective at inhibiting PRV and BoHV-1 entry. MLN4924 inhibited only ∼23% of PRV BeBlue entry at the highest concentration tested (Fig. 3C). MLN4924 inhibited only ∼44% of BoHV-1 v4a entry at the highest concentration tested (Fig. 3D). In contrast, MG132 effectively inhibited both PRV BeBlue and BoHV-1 v4a in a concentration-dependent manner as previously reported (Fig. 3C, D). It is unclear why MLN4924 is less effective at inhibiting PRV and BoHV-1 entry relative to HSV. It may be due to differences in the viruses themselves or in the cell types tested. Concentrations that resulted in ∼50% inhibition of viral entry were: 10 µM MLN4924 and 20 µM MG132 for HSV-1, 1 µM MLN4924 and 5 µM MG132 for HSV-2, 20 µM MG132 for PRV, and 1 µM MG132 for BoHV-1. The concentrations of MLN4924 and MG132 tested had little to no cytotoxicity as measured by LDH assay (Fig 3A-D).

**Figure 3.**
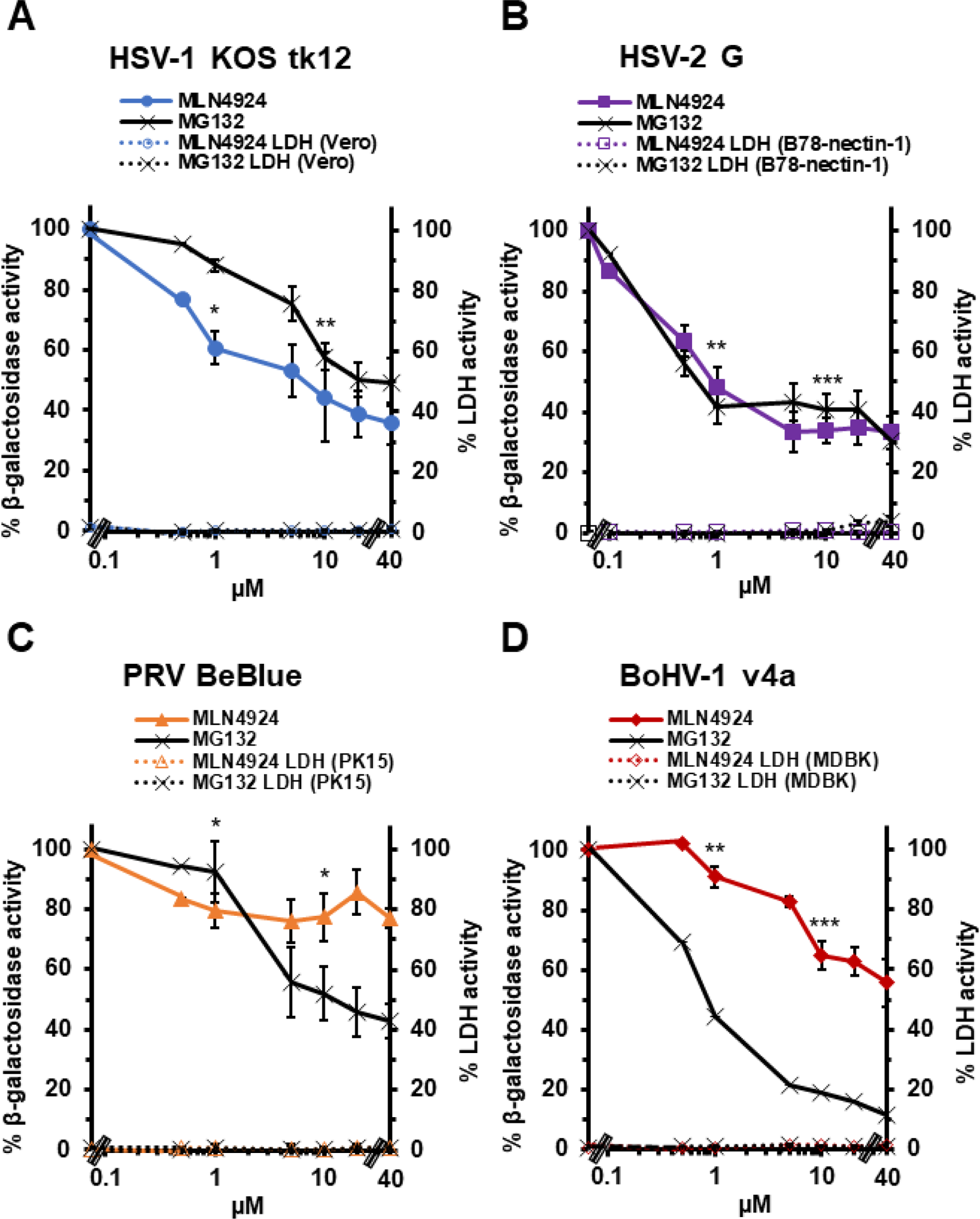
Effect of neddylation inhibitor MLN4924 on alphaherpesvirus entry. Vero cells (A), B78-nectin-1 cells (B), PK15 cells (C), or MDBK cells (D) were treated with MLN4924 or MG132 for 20 min at 37°C. HSV-1 KOS tk12 (MOI of 6) (A), HSV-2 G (MOI of 0.1) (B), PRV BeBlue (MOI of 1) (C), and BoHV-1 v4a (MOI of 2) (D) was added to cells in the continued presence of drugs. At 6 h p.i., cell lysates were prepared, and β-galactosidase activity of the mock-treated, infected cells was set to 100%. Cytotoxicity is shown as percent LDH activity. Values are the means and standard errors of data from three independent experiments. Student’s *t*-test comparing no-drug to 1 µM or 10 µM MLN4924-treated samples. (*, P < 0.05; **, P < 0.01; ***, P < 0.005)

We next ascertained the impact of neddylation on HSV-1 entry into cultured human cells. MLN4924 inhibited HSV-1 entry into human epithelial cells HeLa (Fig. 4A) and HaCaT (Fig. 4B) and into human neuroblastoma cells SK-N-SH (Fig. 4C) and IMR-32 (Fig. 4D). The inhibitor effect was dose-dependent for each cell type, and inhibition was obtained at non-cytotoxic concentrations as assayed by LDH release (Fig. 4). Given the robust effect of MLN4924 on HSV-1 entry (Figs. 3A, 4) and infection (Fig. 2A, D), we moved forward with additional investigation of neddylation and HSV-1 entry.

**Figure 4.**
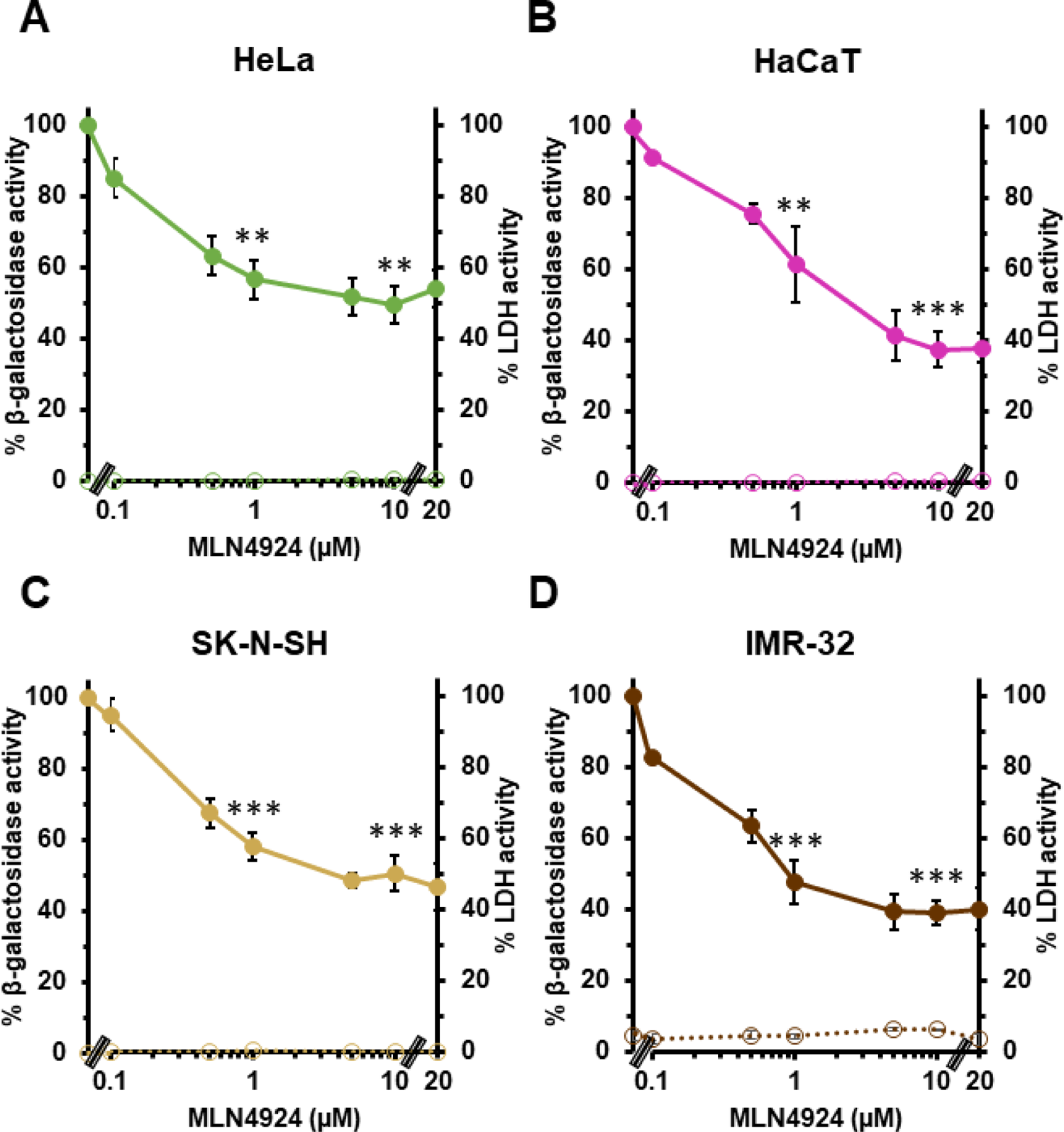
Inhibitory effect of MLN4924 on HSV-1 entry into human cell lines. HeLa (A), HaCaT (B), SK-N-SH (C), or IMR-32 (D) cells were treated with MLN4924 for 20 min at 37°C. HSV-1 KOS tk12 (MOI of 1) was added to cells in the continued presence of drug. At 6 h p.i., cell lysates were prepared, and β-galactosidase activity of the mock-treated, infected cells was set to 100%. Cytotoxicity is shown as percent LDH activity. Concentrations that resulted in ∼50% inhibition of viral entry: 5 µM for HeLa, 1-5 µM for HaCaT, 5 µM for SK-N-SH, and 1 µM for IMR-32. Values are the means and standard errors of data from three independent experiments. Student’s *t*-test comparing no-drug with each indicated MLN4924 treatment. (**, P < 0.01; ***, P < 0.005)

### Role for NEDD8 in HSV-1 entry

More is known about viral entry into model CHO-receptor cells than any other cultured cell type (21-23, 27, 70). MLN4924 treatment of CHO-HVEM cells resulted in inhibition of HSV-1 entry (Fig. 5A). Thus, neddylation functions during HSV entry into all seven cell lines tested in this study. To verify and extend the function of neddylation in HSV entry using a non-pharmacologic approach, we assessed the effect of siRNA knockdown of NEDD8 gene expression. NEDD8 is a ubiquitin-like molecule that is activated and conjugated to substrates in the neddylation cascade (40, 41). CHO-HVEM cells were transfected with siRNA targeting NEDD8 or with a control siRNA. Treatment with the specific siRNA downregulated NEDD8 expression by ∼ 80% as determined by RT-qPCR (Fig. 5B). To determine whether HSV entry requires NEDD8, CHO-HVEM cells were transfected with NEDD8 or control siRNAs and viral entry was measured by the β-galactosidase reporter assay (Fig. 5C). Knockdown of NEDD8 inhibited HSV-1 entry via endocytosis by ∼ 44 % (Fig. 5C). All together the results support the notion that NEDD8 and the NEDD8-activating enzyme facilitate HSV-1 entry.

**Figure 5.**
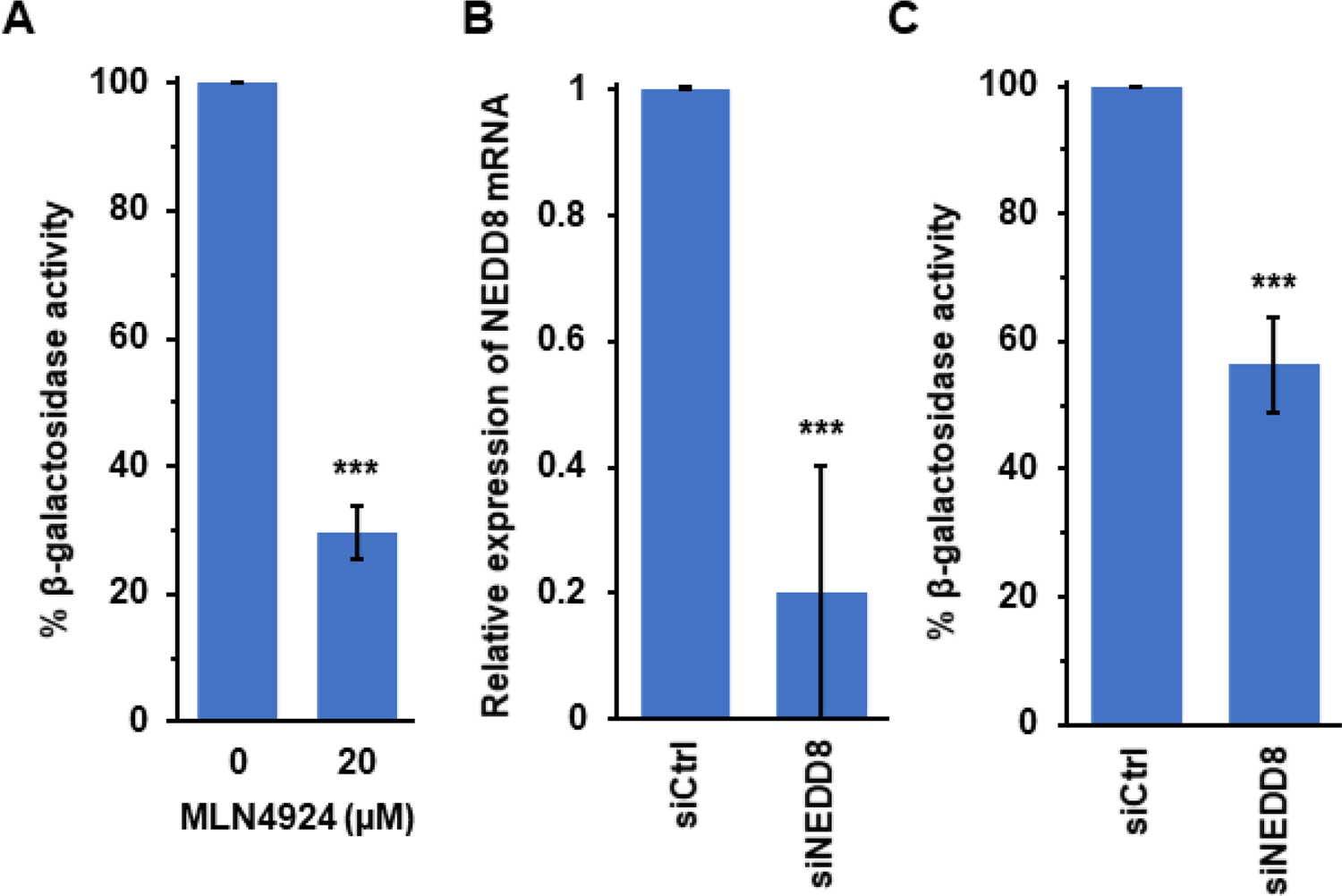
NEDD8 facilitates HSV-1 entry. (A) CHO-HVEM cells were either mock-treated or treated with 20 µM MLN4924 for 20 min at 37°C. HSV-1 KOS (MOI ∼ 0.3) was added. At 6 h p.i., cell lysates were prepared and assayed for β-galactosidase activity as an indicator of viral entry. Activity of mock-treated cells was set to 100%. (B, C) CHO-HVEM cells were transfected with control or NEDD8 siRNAs for 24 h. (B) The efficiency of NEDD8 knockdown was determined by RT-qPCR. Transfected cells were harvested, and total RNA samples were extracted. The level of NEDD8 mRNA expression was determined by the 2^-ΔΔCT^ method using the GAPDH mRNA extracted from control-transfected cells for normalization. (C) The siRNA-treated cells were infected with HSV-1 KOS (MOI ∼ 0.3) and entry was assayed as in (A). Activity of siCtrl-treated cells was set to 100%. Values are the mean and standard deviation of data from three (A, C) or four independent experiments (B). Student’s *t* test (***, *P* < 0.005).

### The inhibitory effect of MLN4924 on HSV entry is reversible and occurs at an early step

Pharmacologic inhibitors of the proteasome can act either in a reversible or irreversible manner (17, 71–73). As an example, MG132 binds reversibly to the active site on the 20S proteasome and has a reversible effect on both proteolytic activity and HSV entry (17, 71). The effect of MLN4924 on NAE activity is partially reversible (56). To further characterize the effect of MLN4924 on HSV-1 entry, we used wash-out experiments to ascertain reversibility. We pre-treated B78-nectin-1 cells with MLN4924 for 20 min and then cultures were either washed to remove the drug or the drug was allowed to remain. HSV-1 was added and entry was measured at 6 h p.i. by β-galactosidase reporter assay (Fig. 6A). Cells that were treated with MLN4924 and then washed prior to addition of virus permitted much more HSV-1 entry than the cells that were not washed. In this system, the inhibitory effect of MLN4924 on HSV entry was 81-93% reversible. This is consistent with the notion that MLN4924 inhibits HSV entry by blocking the enzymatic activity of host cell NAE, since both effects can be reversed by wash-out.

**Figure 6.**
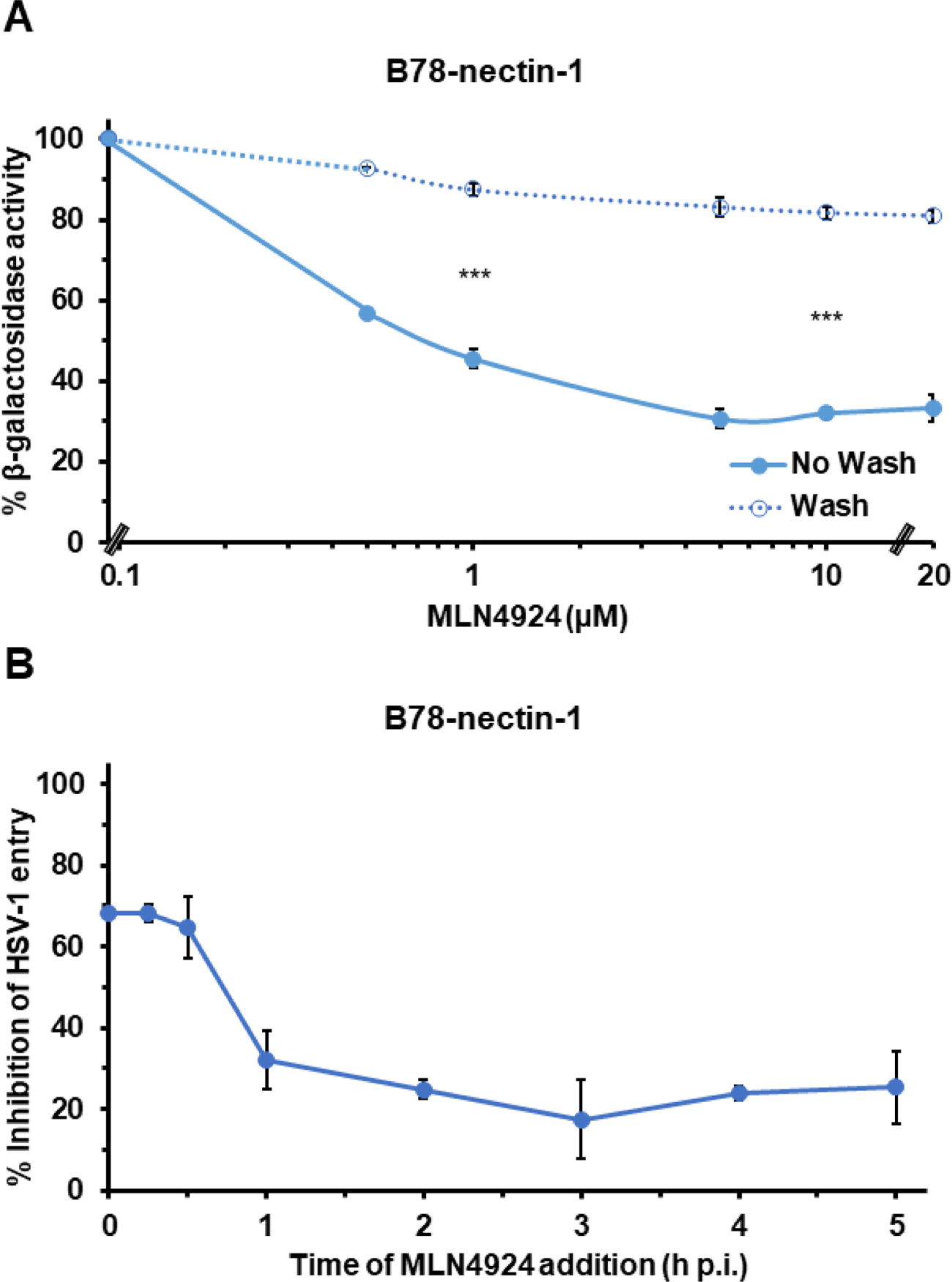
The inhibitory effect of MLN4924 on HSV-1 entry is reversible and early. (A) B78-nectin-1 cells were pretreated with MLN4924 for 20 min at 37°C. HSV-1 KOS (MOI 2) was added in the continued presence of MLN4924 (No Wash), or the medium containing MLN4924 was removed, and the cells were subjected to four 5-min washes with medium prior to adding virus (Wash). At 6 h p.i., cell lysates were prepared, and β-galactosidase activity of mock-treated, infected cells was set to 100%. Student’s *t*-test comparing no-wash with wash at 1 µM or 10 µM MLN4924. (***, P < 0.005) (B) HSV-1 KOS (MOI 2) was bound to B78-nectin-1 cells for 1 h at 4°C. Cells were rapidly warmed to 37°C to initiate infection. At the indicated times, 20 μΜ MLN4924 was added. At 6 hr p.i., cell lysates were prepared and assayed for beta-galactosidase activity. Activity of the mock-treated, infected cells was set to 100% (not shown). Values are the means and standard errors of three independent experiments.

A time-of addition assay was performed to determine whether MLN4924 inhibits HSV-1 at an earlier or later step of infection. HSV-1 KOS was bound to B78-nectin-1 cells at 4°C for 1 h, and cultures were then shifted to 37°C. At 0 to 6 h p.i., 20 µM MLN4924 was added. The later the treatment, the less of an inhibitory effect on HSV entry was detected. This suggested that MLN4924 acts on an early step in HSV infection (Fig. 6B). Together, the results suggest that neddylation mediates an early, postbinding step in HSV entry, such as endocytic uptake from the B78-nectin-1 cell surface.

### Inhibition of neddylation blocks HSV entry at the level of endocytic internalization from the plasma membrane

HSV-1 entry can be serialized into several sequential steps. Following attachment to the cell surface, HSV-1 is either internalized by an endocytosis mechanism or directly penetrates into the cytosol (29, 74–76). B78-nectin-1 and HaCaT cells support HSV endocytosis, and Vero cells support direct penetration (fusion) of HSV at the plasma membrane. To understand better the specific HSV entry step that is neddylation-dependent, we measured the effect of MLN4924 on the uptake of infectious HSV into the different cells (Fig. 7). HSV-1 was first bound to cells at 4°C. At various times after warming of cultures to 37°C, extracellular virus was inactivated. Cultures were incubated for a total of 18-24 h, and plaque formation was determined. The acquisition of infectious virion resistance to inactivation reflects internalization by endocytosis (B78-nectin-1 or HaCaT cells) or fusion with the plasma membrane (Vero cells). HSV-1 internalization into mock-treated B78-nectin-1 cells occurred rapidly with a t_1/2_ of ∼ 10 min (Fig. 7A). MLN4924-treatment inhibited HSV-1 internalization into B78-nectin-1 cells by > 90% (Fig. 7A). Furthermore, MLN4924-treatment of the human epidermal keratinocyte line HaCaT inhibited virus internalization by ∼84% (Fig. 7B). To probe further the striking role of neddylation in endocytic internalization of HSV-1, we determined the contribution of proteasome activity to HSV-1 internalization in B78-nectin-1 cells (Fig. 7C). MG132 had a more modest effect on the overall endocytic internalization of HSV-1 (∼38% inhibition; Fig. 7C) than did MLN4924. Further, MG132 had little effect on the kinetics of HSV internalization. The rate at which HSV-1 was internalized in the presence of MG132 was similar to mock-treated (Fig. 7C).

**Figure 7.**
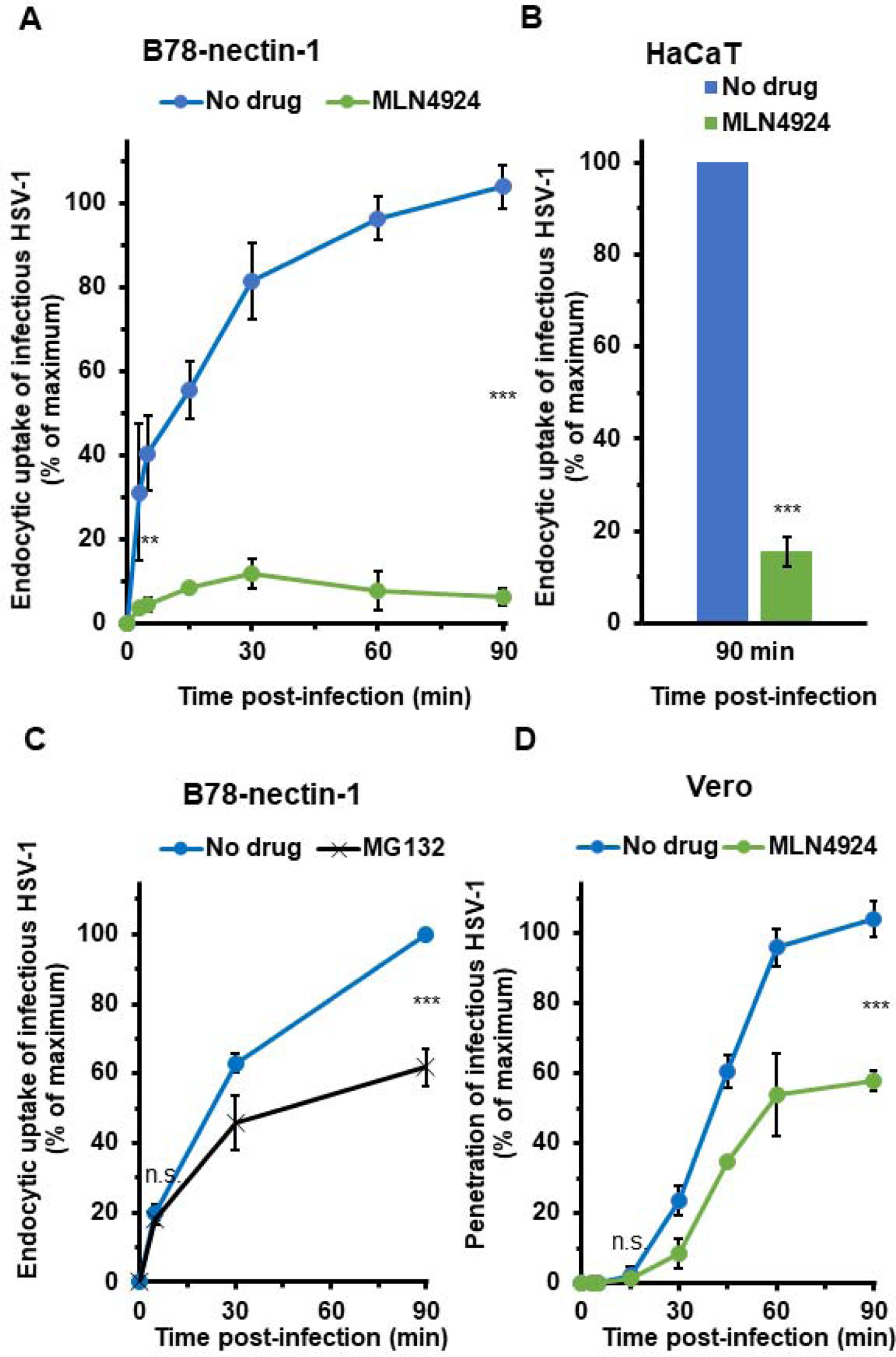
Neddylation mediates uptake of HSV-1 from the cell surface. B78-nectin-1 (A, B), HaCaT (B), or Vero (C) cells were pre-treated with 20 µM MLN4924 (A, B, D) or MG132 (C) for 20 min at 37°C. Cells were chilled for 10 min at 4°C on ice. HSV-1 KOS (approximately 100 PFU/well) was added for 1 h at 4°C on ice in the continued presence of drug to allow for virus attachment. Cells were washed with ice-cold PBS and incubated at 37°C. At the indicated times p.i., extracellular HSV-1 was inactivated with sodium citrate buffer (pH 3.0). At 18-24 h p.i., cells were fixed, and plaque formation was determined. Average plaque numbers at 60-90 min p.i. was set to 100%. Values are the mean and standard deviation of data from three independent experiments. Student’s *t*-test comparing no-drug with MLN4924-treated at 5 min or 90 min p.i. (**, P < 0.01; ***, P < 0.005)

We assessed the role of neddylation in HSV fusion with the plasma membrane. MLN4924-treatment of Vero cells inhibited direct penetration of HSV-1 by ∼42% (Fig. 7D). The kinetics of HSV that successful entered Vero cells via virus-cell fusion in the presence of MLN4924 was similar to that of mock-treated cells (Fig. 7D). These data suggest that neddylation plays a major function in HSV internalization, but it may also impact direct penetration. We have shown previously that MG132 does not directly inhibit virus-Vero cell fusion during entry nor virus-induced fusion of Vero cells (17). The present results suggest that HSV-1 endocytic internalization is less reliant on proteasome activity than it is on neddylation. It is of future interest to test the effect of combining inhibitors of neddylation and proteasomal degradation on virus internalization. Monoubiquitination mediates membrane traffic independent of proteasome activity (77–79). It is tempting to speculate that neddylation similarly affects endocytic internalization and that HSV is capitalizing on such a process for its entry. All together, our findings suggest that host cell neddylation activity is critically important for surface uptake of HSV-1 by endocytosis.

## DISCUSSION

Proteasome activity assists the entry of many viruses, including the alphaherpesviruses. We report that the proteasome inhibitor bortezomib impairs entry and infection by HSV-2, PRV, and BoHV-1, consistent with a conserved proteasome-dependent mechanism employed by alphaherpesviruses (17–20). The host cell neddylation pathway is important for cell infection by all alphaherpesviruses tested. However, the NEDD8 pathway plays a specific role in entry of HSV-1 and HSV-2, but not PRV or BoHV-1. We propose that endocytic uptake of entering HSV-1 virions from the plasma membrane is a neddylation-dependent process. This is the first report of neddylation mediating virus entry to our knowledge.

HSV-1 tegument ICP0 regulates the proteasome-dependent delivery of entering HSV-1 capsids to the host cell nucleus (17, 20, 30, 31). The involvement of ICP0 orthologs in the entry mechanism of other alphaherpesviruses has yet to be investigated. The nature and identity of the substrate targeted for proteasomal degradation during alphaherpesvirus entry also remains to be determined. The peptide boronate bortezomib is a cancer drug, and was the first proteasome inhibitor to be FDA-approved for clinical use (52, 53, 80–82). The potential repurposing of bortezomib as an anti-HSV agent has been discussed in detail (20). We provide evidence that bortezomib is a pan-alphaherpesvirus entry inhibitor, blocking the entry of PRV, BoHV-1 and HSV-2, in addition to HSV-1.

NEDD8 is a 9 kDa, 81 amino acid ubiquitin-like molecule (40). Neddylation is a post-translational modification that conjugates NEDD8 to a lysine residue of a target protein. The neddylation pathway is analogous to ubiquitination but uses its own E1 activating and E2 conjugating enzymes (41, 42). The largest family of E3 ubiquitin ligases are the cullin-RING ligases (CRLs), which are activated by NEDD8, resulting in ubiquitination of substrate proteins (83–86). CRL modification is the most well-studied neddylation process and illustrates a direct link between neddylation and the UPS (43, 87–89). The proteasome-dependent entry of HSV-1 entry does not require activation of ubiquitin (17). Therefore, we assessed whether neddylation aided in the entry of alphaherpesviruses.

MLN4924 is a cell-permeable, adenosine monophosphate mimetic that binds to NAE, blocking the NEDD8 cascade. It is being tested as a cancer drug in Phase 1 and Phase 2 clinical trials (57, 58, 90, 91). MLN4924 blocks infection by the alphaherpesviruses tested, consistent with reports that MLN4924 inhibits infection by many viruses (38, 39, 46, 49–51). The antiviral activity of MLN4924 may involve NEDD8-conjugated CRLs (38, 39). We show that MLN4924 inhibits HSV-1 and HSV-2 in viral entry assays. MLN4924 has less of an inhibitory effect on the entry of PRV and BoHV-1 than it does on infection by these veterinary viruses, highlighting that post-entry functions in the alphaherpesvirus replication cycles are likely reliant on neddylation. Our experimental results indicate that MLN4924’s effect on HSV entry can be reversed, consistent with the reversible effect of MLN4924 on the ability of NAE to conjugate NEDD8 to substrates.

There are few if any host cell processes known to mediate the initial internalization step of HSV entry. Endocytic uptake of HSV can occur independently of clathrin, caveolin, cholesterol, and dynamin (22, 68, 92–97). We show that neddylation is integral to internalization of HSV-1 by endocytosis in B78-nectin-1 cells and human HaCaT cells. HSV-1 internalization in CHO cells is thought to be independent of nectin-1 and other gD-receptors (70). However, in B78 mouse melanoma cells, nectin-1 mediates HSV-1 uptake (21). The interaction of HSV-1 gH with integrins is thought to facilitate viral internalization (98). HSV-1 envelope proteins gC, gE, gG, gI, gJ, gM, UL45, and US9 are dispensable for entry by endocytosis (60, 99, 100), but the impact of these proteins specifically on internalization is not clear. The E3 ubiquitin ligase Cbl together with HSV-1 ICP0 contributes to the downregulation of surface-expressed nectin-1 at late stages of HSV-1 infection; however, depletion of Cbl has no inhibitory effect on viral entry (101). Additionally, our results suggest that while neddylation mediates endocytic uptake of HSV-1, it likely also contributes to other steps in HSV entry and infection, which have yet to be investigated. For example, MLN4924 negatively affects virus-cell fusion, although not as dramatically as it inhibits internalization (>90% inhibition) (Fig. 7).

Ubiquitination of plasma membrane proteins is frequently sufficient for their endocytic internalization, independent of proteasomal degradation (102–104). Interplay between NEDD8-modification and Ub-modification of substrate proteins leads to their endocytic trafficking, and influences endocytosis signaling pathways (105, 106). The substrate that is tagged by NEDD8 during HSV internalization remains an important question. In summary, we propose that the initial step of HSV entry by endocytosis is mediated by the neddylation pathway.

## MATERIALS AND METHODS

### Cells

B78 murine melanoma cells expressing nectin-1 (B78-nectin-1 or C10 cells) (107), a gift from G. Cohen and R. Eisenberg (University of Pennsylvania), were propagated in Dulbecco’s modified Eagle medium (DMEM; Thermo Fisher Scientific, Waltham, MA) supplemented with 10% fetal bovine serum (FBS; Atlanta Biologicals. Atlanta, GA) and 1% penicillin, streptomycin, and glutamine (PSG, Thermo Fisher Scientific). Selection of B78-nectin-1 occurred every fourth passage in medium further supplemented with 250 μg/ml of Geneticin (Sigma, St. Louis, MO) and 6 μg/ml of puromycin (Sigma). CHO-HVEM (M1A) cells, a gift from R. Eisenberg and G. Cohen were propagated in Ham’s F12 nutrient mixture (Gibco/Life Technologies) supplemented with 10% FBS and 1% PSG. Selection of CHO-HVEM cells occurred every fourth passage in medium supplemented with 150 μg/ml of puromycin and 250 μg/ml of G418 sulfate (Thermo Fisher Scientific, Fair Lawn, NJ). Both B78-nectin-1 and CHO-HVEM cells contain the *Escherichia coli lacZ* gene under the control of the HSV-1 ICP4 promoter. PK15 cells, a gift from Matthew Taylor (Montana State University) and Vero cells (American Type Culture Collection (ATCC), Manassas, VA) were propagated in DMEM supplemented with 10% FBS and 1% PSG. Madin Darby bovine kidney (MDBK) cells (ATCC) were propagated in DMEM supplemented with 5% FBS and 1% PSG. Human HaCaT epithelial keratinocytes and HeLa cells were propagated in DMEM supplemented with 10% FBS. Nondifferentiated human SK-N-SH and IMR-32 neuroblastoma cells (ATCC) were propagated in Eagle’s minimal essential medium supplemented with 10% FBS, 1 mM sodium pyruvate, 0.1 mM nonessential amino acids, and Earle’s salts (Invitrogen).

### Viruses

HSV-1 strain KOS, a gift from Priscilla Schaffer (Harvard Medical School), was propagated and titered on Vero cells. HSV-1 strain KOS tk12, a gift from Patricia Spear (Northwestern University), contains the *Escherichia coli lacZ* gene inserted into the thymidine kinase gene under the control of the HSV-1 infected-cell protein 4 (ICP4) promoter. The ICP4 gene is expressed with immediate-early kinetics. HSV-2 strain G was obtained from ATCC. All HSV-1 and HSV-2 viruses were propagated on Vero cells. PRV BeBlue, a gift from Lynn Enquist (Princeton University), is a recombinant PRV Becker strain with the *Escherichia coli lacZ* gene inserted into the gG locus (108). The gG gene is expressed with early kinetics. PRV BeBlue was propagated and titered on PK15 cells. BoHV-1 v4a, a gift from J. C. Whitbeck, G. Cohen, and R. Eisenberg (University of Pennsylvania), is a recombinant of BoHV-1 Colorado-1 strain and contains the *Escherichia coli lacZ* gene in place of the viral thymidine kinase (TK) gene (109). The TK gene is expressed with early kinetics. BoHV-1 v4a was propagated and titered on MDBK cells. In the reporter assay for entry, the infectivity of all *lacZ^+^* viruses was measured at 6 h p.i.

### Chemicals

Stocks of 50 mM bortezomib (Selleckchem, Houston, TX, or Sigma, St. Louis, MO) were prepared in dimethyl sulfoxide (DMSO; Fisher Scientific, Fair Lawn, NJ) and stored at −80°C. Stocks of 20 mM MG132 (Sigma) were prepared in DMSO and stored at −20°C. Stocks of 20 mM MLN4924 from BostonBiochem (Cambridge, MA) or Tocris (Minneapolis, MN) were stored in DMSO at −20°C. Drugs were diluted in the appropriate cell culture medium to achieve 0 to 40 μM concentrations immediately prior to use.

### Βeta-galactosidase reporter assay of viral entry

Confluent monolayers of cells were infected with HSV-2 G (MOI 0.1), HSV-1 KOS tk12 (MOI 6), HSV-1 KOS (MOI 0.3 or 2), PRV BeBlue (MOI of 1), or BoHV-1 v4a (MOI of 2) at 37°C. At 6 h p.i., cells were lysed with 0.5% IGEPAL (Sigma–Aldrich) followed by one freeze-thaw cycle. Chlorophenol red-beta-D-galactopyranoside (Roche Diagnostics, Indianapolis, IN) substrate was added to cell lysates, and the β-galactosidase activity was read at 595 nm with an ELx808 microtiter plate reader (BioTek Instruments, Winooski, VT).

### Determination of cytotoxicity of pharmacologic agents

Cells were treated with drugs under experimental conditions in the absence of virus. Cytotoxicity was quantified by direct measurement of lactate dehydrogenase (LDH) with a colorimetric assay (110). LDH leakage was determined using the CyQUANT LDH Cytotoxicity Assay Kit (Invitrogen) according to the manufacturer’s instructions. Cell viability was calculated by setting mock-treated cells to 100% and subtracting the cytotoxicity value.

### Alphaherpesvirus plaque assays

Sub-confluent cell monolayers were grown in 24-well plates. For plaque formation experiments, 80-120 PFU of virus per well was added to cells. Cultures were incubated for 1 – 1.5 h at 37°C followed by addition of warm DMEM with 5% fetal bovine serum and 2% carboxymethyl cellulose (CMC; Sigma) to achieve a 1% CMC overlay. At 44 - 48 h p.i., cells were fixed with 10% formalin (VWR International, Solon, OH) and stained with crystal violet (Sigma). Plaques that excluded stain were enumerated. For infectivity experiments, MOIs were approximately 0.01 for HSV-1, 0.0002 for PRV, and 0.002 for BoHV-1. After 24 h p.i., cells were scraped and combined with supernatant. After a freeze-thaw cycle and sonication, the lysate was titered by limiting dilution, crystal violet plaque assay as described above.

### NEDD8 knockdown with siRNA

CHO-HVEM cells were grown in 24-well plates to 70% confluence. Culture medium was removed, and the cells were transfected with siCtrl (sc-37007) or siNEDD8 (sc-36026) (Santa Cruz Biotechnology) using Lipo3000 transfection reagent (Thermo Fisher). Briefly, 60 pmol of siRNA were mixed with 1.5 μl Lipo3000 in 50 μl of OPTIMEM (Gibco Life Technologies) without serum for 15 min at room temperature. Serum-free OPTIMEM medium was added to a final volume of 250 μl, and the transfection reaction was added to the cells for 5 h at 37°C. The transfection mixture was replaced with medium supplemented with 10% FBS, and cultures were incubated for 24 h at 37°C prior to infection with HSV-1.

### Quantification of NEDD8 knockdown by qPCR

Approximately 1 × 10^5^ CHO-HVEM cells were plated in a 24-well plate and the subsequent day transfected with siCtrl (Santa Cruz Biotechnology) or siNEDD8 (Santa Cruz Biotechnology) as previously described (27). At 24 h post-transfection, total RNA was isolated with TRIzol reagent (Invitrogen) and chloroform (Baker) according to the manufacturer’s instructions. Finally, RNA was resuspended in 25 μl of DEPC-treated water (Ambion). Contaminating DNA was removed with TURBO DNA-free kit (Thermo) and the absence of DNA was confirmed by qPCR. Primers were purchased from Integrated DNA Technologies (Coralville, Iowa): Forward sense NEDD8 primer 5’-AAG GTG GAG CGA ATC AAG GA-3’; reverse antisense NEDD8 primer 5’-GCT TGC CAC TGT AGA TGA GC-3’; forward sense GAPDH primer 5’-CGA CTT CAA CAG CAA CTC CCA CCT CTT CC-3’; reverse antisense GAPDH primer 5’-TGG GTG GTC CAG GGT TTC TTA CTC CTT-3’. RNA concentration was determined by absorbance at 260 nm with a NanoDrop spectrophotometer (DeNovix DS-11 Series). 1 μg of RNA from each sample was reverse-transcribed into cDNA for qPCR analysis. Reverse transcription was performed with SuperScript VILO cDNA Synthesis Kit (Thermo Fisher) in a final volume of 20 μl. cDNAs were diluted 1:50 with DEPC-treated water (Ambion), then subjected to real-time qPCR analysis. Real-time quantitative PCR experiments were performed with BioRad CFX96 Real Time System. All reactions were carried out in 20-μl reaction mixtures using SsoAdvanced Universal SYBR Green Supermix (BioRad). Relative expression levels were calculated by the 2^-ΔΔCT^ method (111) with GAPDH as the internal reference.

### Cell type-dependent endocytic internalization or direct penetration of HSV-1

B78-nectin-1 cells (for internalization) or Vero cells (for penetration) were pre-treated with MLN4924 or MG132 diluted in DMEM for 20 min at 37°C. Medium was replaced with ice-cold, carbonate-free, serum-free DMEM supplemented with 20 mM HEPES and 0.2% bovine serum albumin (binding medium). Cells were chilled for 10 min at 4°C on ice. HSV-1 KOS was added (∼100 PFU/well) in ice-cold binding medium containing drug for 1 h at 4°C on ice. Cells were rinsed with ice-cold PBS and warm complete DMEM was added. Cultures were returned to 37°C. At the indicated times p.i., cells were treated with warm sodium citrate buffer (pH 3.0) for 3 min at 37°C to irreversibly inactivate extracellular virions (112). Complete DMEM was added and the infection was allowed to continue at 37°C. At 18-24 h p.i., cells were fixed with methanol-acetone (2:1). Cells were stained with rabbit polyclonal antibody to HSV (HR50; Fitzgerald Industries, Acton, MA) followed by a horseradish peroxidase-conjugated protein A (Thermo Fisher). Finally, 4-Choloro-1-naphthol substrate (Sigma) and H_2_O_2_ catalyst (VWR International, Inc., Radnor, PA) were added to visualize plaques.

## ACKNOWLEDGMENTS

We thank Gary Cohen, Roselyn Eisenberg, Lynn Enquist, Matthew Taylor, J. Charles Whitbeck and Patricia Spear for providing reagents. We thank Reginaldo Bastos for helpful discussions. This study was supported by National Institutes of Health grants R01 AI119159 (A.V.N) and T32 AI007025 (B.H.L.).

